# Dynamical instability measured by temporal entropy improves psychiatric classification across cohorts

**DOI:** 10.64898/2026.04.28.721265

**Authors:** Tatsuma Shoji, Ryo Nakaki

## Abstract

Psychiatric disorders, such as attention-deficit/hyperactivity disorder, autism spectrum disorder, and schizophrenia, are clinically heterogeneous and lack objective biomarkers for reliable diagnosis. Although blood transcriptomic data have been proposed as a potential source of diagnostic information, their generalizability across independent cohorts remains unclear. This study aimed to assess whether biologically informed measures of dynamic instability enhance the reproducibility and generalizability of psychiatric classifications based on peripheral blood data by integrating publicly available blood transcriptomic datasets from multiple cohorts and evaluating classification performance using individual-level cross-validation and study-level holdout validation. To investigate the underlying biological structure, we applied a dynamic systems framework, including pseudotime-based vector field inference and attractor analysis. Additionally, we introduced temporal entropy as a measure of dynamic instability in the inferred transcriptomic trajectories. High classification performance was observed in individual-level cross-validation (area under the receiver operating characteristic [AUROC] > 0.8 across several comparisons); however, performance decreased substantially in study-level validation (AUROC ≈ 0.5–0.7), indicating limited generalizability. Attractor analysis revealed that transcriptomic states formed continuous and overlapping structures rather than distinct diagnostic clusters. Stratification based on temporal entropy identified a subset of individuals with unstable transcriptomic dynamics, and excluding these individuals improved the classification performance across most diagnostic pairs (AUROC > 0.7). These findings suggest that transcriptomic variability and dynamic instability contribute to the limited reproducibility of psychiatric classifications. Incorporating temporal entropy as a measure of system-level instability may enhance the robustness and interpretability of biomarker-based models and provide a new perspective on psychiatric disorders as dynamic systems.

## 1 Background

Psychiatric and neurodevelopmental disorders, including attention deficit hyperactivity disorder (ADHD), autism spectrum disorder (ASD), and schizophrenia (SCZ), present significant global public health challenges due to their substantial impact on individual functioning, family burden, and socioeconomic systems. The prevalence of these conditions is notable, with ADHD affecting approximately 5% of children worldwide (1) and ASD occurring in approximately 1 in 100 individuals globally (2). Although SCZ is less prevalent, it disproportionately contributes to the global disease burden because of its severity (3). Early identification and intervention have improved developmental and functional outcomes, as shown in randomized trials of early intervention for ASD (4) and emphasized in clinical guidelines for ADHD and early psychosis (5,6). However, current diagnostic practices rely primarily on clinical interviews and behavioral assessments, which are subjective and susceptible to variability in symptom presentation, comorbidities, and environmental factors. Consequently, a growing need exists for objective, biologically grounded biomarkers that complement clinical evaluation and enhance diagnostic precision.

Despite the clinical importance of early and accurate diagnosis, current psychiatric classification heavily depends on subjective assessment, leading to diagnostic uncertainty and delays, particularly in ASD, where no established laboratory biomarkers exist (7). To address this limitation, interest has increased in developing objective biomarkers based on peripheral biological signals, including blood transcriptomic profiles. Peripheral blood is particularly attractive because of its accessibility and partial correspondence between the blood and brain molecular signatures. The potential of blood-based gene expression signatures for classifying psychiatric conditions, such as ASD, ADHD, and SCZ (8). However, a critical limitation of these approaches is their lack of generalizability across independent cohorts. High-throughput omics data are susceptible to batch effects and study-specific confounding factors, including population characteristics, technical platforms and experimental conditions (9). When such confounding factors are correlated with diagnostic labels, predictive models may learn study-specific artifacts rather than disease-related signals, leading to overestimated performance (10). Therefore, robust biomarker development in multi-cohort settings requires appropriate batch correction methods, such as ComBat (11), and rigorous evaluation frameworks that assess generalization across independent studies (12,13).

Beyond static diagnostic categories, psychiatric disorders can be viewed as dynamic systems, characterized by temporal fluctuations in symptoms and underlying biological states. Theoretical frameworks from systems biology, such as attractor dynamics and Waddington’s epigenetic landscape, suggest that cellular states may transition between multiple stable and quasi-stable configurations (14,15). In parallel, studies on psychological time-series data have proposed early warning signals for state transitions, including increased variability and critical slowing, prior to abrupt changes in mental states (16). These perspectives motivate the use of dynamic features derived from cross-sectional data as proxies for underlying temporal instabilities.

In this study, we integrated publicly available blood transcriptomic datasets across multiple cohorts and evaluated a unified analytical framework consisting of (i) conventional pairwise classification based on individual-level random splits; (ii) more stringent evaluation using study-level holdout validation to assess generalizability; (iii) inference of transcriptomic dynamics via pseudotime-based vector field modeling and attractor analysis; and (iv) stratification of individuals based on temporal entropy, a metric designed to quantify the instability of inferred transcriptomic trajectories. By combining these approaches, we aimed to determine whether biologically informed measures of dynamic instability can improve the reproducibility and generalizability of psychiatric classifications based on peripheral blood data.

## 2 Methods

### 2.1 Study Design and Reporting Framework

A secondary analysis of publicly available transcriptomic datasets was conducted to develop and evaluate predictive models for psychiatric classification based on peripheral blood gene expression. The analytical framework integrates machine learning classification with dynamic system analysis to assess the predictive performance and underlying structure of transcriptomic variability. To evaluate model performance under varying assumptions of data independence, we used two complementary validation strategies: (i) individual-level cross-validation with random sample partitioning and (ii) study-level holdout validation, where the entire cohort was excluded from training and used exclusively for testing. This design facilitated the assessment of within-cohort predictive performance and cross-cohort generalizability. All analyses were conducted retrospectively using de-identified, publicly accessible data; thus, additional ethical approval was not required.

### 2.2 Data Sources and Cohort Description

Publicly available blood transcriptomic datasets were collected for three psychiatric conditions: ADHD, ASD, and SCZ, along with healthy controls (CTRL). Seven independent cohorts were included in this analysis. For ADHD, RNA sequencing data from the Gene Expression Omnibus (GEO) dataset GSE159104 comprised two cohorts: a twin cohort (16 CTRL and 16 ADHD) and non-twin cohort (21 CTRL and 23 ADHD). For ASD, RNA sequencing data from GSE212645 (53 CTRL and 51 ASD) and microarray data from GSE18123 (8) included two independent cohorts (82 CTRL, 41 ASD, 21 CTRL, and 21 ASD). For SCZ, microarray datasets GSE38484 (17) (96 CTRL, 106 SCZ) and GSE38481 (17) (22 CTRL, 15 SCZ) were used. The integrated dataset encompassed 584 samples from all cohorts, including individuals with ADHD, ASD, SCZ, and CTRL. These datasets represent a heterogeneous collection of transcriptomic profiles generated using various platforms and study designs, providing a suitable basis for evaluating cross-cohort generalizability.

### 2.3 Data Preprocessing and Integration

Raw transcriptomic data from each cohort were independently processed before integration. For RNA sequencing datasets, gene expression values were aggregated at the gene level by averaging replicate measurements and collapsing splice variants into single-gene features. The expression values were converted to transcripts per million, then log2-transformed, and standardized. For microarray datasets, expression values were standardized within each cohort. After preprocessing, genes common to all datasets were retained, resulting in a shared feature set of 6,369 genes. Batch correction was performed using the ComBat method from the Scanpy Library (version 1.9.3) to mitigate effects arising from differences in experimental platforms, sample processing, and study-specific factors. Batch labels corresponding to cohort identity were used for correction. Following batch correction, all cohorts were integrated into a single dataset for downstream analysis.

### 2.4 Pairwise Classification with Stratified Cross-Validation

For pairwise classification, samples from each diagnostic group (CTRL, ADHD, ASD, and SCZ) were extracted from the integrated dataset and encoded as binary outcome variables. The dataset was randomly partitioned into five folds while preserving class proportions (stratified sampling). In each iteration, four folds were used for training and one-fold for testing. This process was repeated five times, ensuring that each fold served as the test set exactly once, yielding out-of-fold (OOF) predictions for all samples. Within each training set, gene expression features (6,369 genes) were standardized using parameters derived exclusively from the training data, with the same transformation applied to the corresponding test data to prevent data leakage. Three machine-learning models were trained: elastic net-regularized logistic regression, linear support vector machine (SVM), and random forest. Hyperparameters for each model were optimized using five-fold stratified cross-validation within the training data. All models were implemented using the Scikit-learn library (version 1.3.0). Final predictions for each sample were obtained from the corresponding OOF test predictions, ensuring that all reported performance metrics were based on out-of-sample evaluations.

### 2.5 Study-Holdout Cross-Validation for Generalization Assessment

We implemented a study-holdout cross-validation framework to evaluate the generalizability of the model and minimize potential study-specific confounding factors. In this framework, train–test splits were defined at the cohort level rather than at the individual level. In each iteration, an entire cohort served as the test set, whereas the remaining cohorts were used for model training. The same pairwise classification tasks, feature sets, preprocessing procedures, and machine learning models were applied in this framework. Within each training set, gene expression features were standardized using parameters derived exclusively from the training data, with the same transformation applied to this held-out cohort. Hyperparameter tuning was conducted using stratified cross-validation within the training cohorts. This procedure was repeated for all possible cohort–holdout combinations to ensure that each cohort served as an independent test set. Final predictions for each sample were derived from the corresponding held-out evaluation, providing a rigorous assessment of cross-cohort generalization performance.

### 2.6 Receiver Operating Characteristic Analysis

The model performance was evaluated using receiver operating characteristic (ROC) analysis based on OOF predicted probabilities. For each classification task and validation framework, predictions across all folds were aggregated so that each sample contributed to a single prediction during out-of-sample evaluation. The central ROC curve was constructed using pooled OOF predictions by computing the true positive rate (TPR) and false positive rate (FPR) across all samples. To assess variability across cross-validation folds, ROC curves were independently computed for every fold. Each fold-specific ROC curve was interpolated onto a common grid of FPR values, and the standard deviation of the TPR values across the folds was calculated at each point. This variability was visualized as a confidence band around the mean ROC curve, defined as mean TPR ± standard deviation, with values constrained to the range [0, 1]. The area under the ROC curve (AUROC) with standard deviation was calculated as the primary performance metric.

### 2.7 Inference of Transcriptomic Dynamics and Attractor Analysis

To investigate the dynamic structure of transcriptomic states from cross-sectional data, we inferred a continuous dynamic system representing gene expression changes along a pseudotemporal trajectory. For each sample, a gene expression state vector *X* was constructed using 300 of the variable genes from the integrated dataset. Selecting the 300 most variable genes captures the dominant sources of transcriptomic variation while reducing noise and computational complexity. A one-dimensional pseudo-time was assigned to each sample based on the first principal component (PC1) of the standardized gene expression matrix. Principal component analysis (PCA) was applied, and PC1 scores were linearly rescaled to [0, 1] to define the pseudotemporal ordering of samples. Gene expression dynamics along pseudotime were modeled using univariate smoothing splines. For each gene *g*, a smooth function *X*_*g*_ (*t*) was fitted. Both the smoothed expression values and their first derivatives, *dX*_*g*_ /*dt* were evaluated at the pseudotime of each sample. These derivatives were interpreted as approximations of the transcriptomic velocity. To model the global dynamics of the system, derivatives *dX*/*dt* were regressed on gene expression state vectors *X* using Ridge regression. This approach yields an approximate deterministic vector field of the form *dX*/*dt* =*F*(*X*), where *F*(*X*) represents the inferred dynamics of the transcriptomic system. Based on this model, the predicted velocity vectors and their Euclidean norms ║*F*(*X*) ║ were computed for all samples. The samples with small velocity norms were interpreted as having dynamically slow or near-stationary states. Accordingly, quasi-fixed points were defined as samples below the 2% lower quantile of the velocity-norm distribution. To further identify the putative exact stationary states, numerical root-finding was performed using quasi-fixed points as initial conditions. For each seed state *X*_0_, a nonlinear solver was used to find solutions of *F*(*X*)= 0, and solutions were retained as true fixed points if the residual norm ║*F*(*X*) ║ was below 10^-6^. Nearby solutions were merged based on the Euclidean distance in the gene-expression space. To evaluate the local dynamic stability, the Jacobian matrix of the inferred vector field was estimated numerically at each quasi-fixed point, and the fixed point was identified using central finite differences. The eigenvalues of the Jacobian were computed, and a state was classified as an attractor when the largest real part of the eigenvalue was negative, indicating local asymptotic stability. The pseudotime variable, denoted as t, does not represent physical time but rather an inferred ordering along the dominant axis of transcriptomic variation. Therefore, the inferred dynamics reflect progression along a latent trajectory rather than directly observed temporal processes.

### 2.8 Definition of Temporal Entropy

To quantify the temporal variability of transcriptomic dynamics, we defined a metric, temporal entropy, which captures the directional and magnitude variability of state transitions along simulated trajectories. Starting from the inferred dynamical system *dX*/*dt = F*(*X*), each sample was propagated forward using numerical integration with the fourth-order Runge–Kutta method for 100 steps, yielding a discrete trajectory *X*(*t*) for each sample. At each step, the instantaneous state change (velocity) is defined as

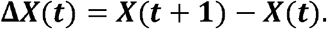

For each sample, a rolling window of the most recent state changes (*W* = 8) was maintained to capture the local trajectory behavior. Temporal entropy was computed as a weighted combination of the directional and speed variabilities within this window. The directional variability was quantified by first normalizing each state-change vector and then computing the magnitude of the mean direction.

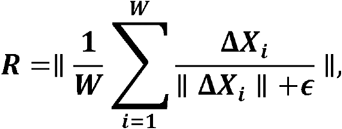

where *ϵ* is a small constant introduced for numerical stability (*ϵ* = 10^−8^). The directional entropy is then defined as

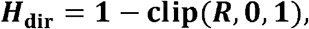

such that consistent directional motion yields low entropy, whereas incoherent motion yields high entropy. Speed variability was computed from the magnitude of state changes as follows:

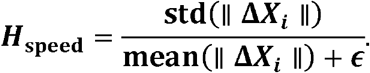

To constrain this term to a bounded range, it is transformed using a saturating function:

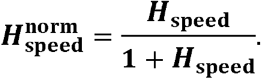

The final temporal entropy was defined as

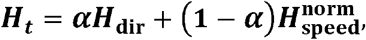

where *α* = 0.7controls the relative contribution of directional versus speed variability. The resulting entropy values were clipped to a range [0’1]. Temporal entropy reflects variability along simulated trajectories from an inferred dynamic system and should be interpreted as a model-based measure of dynamical heterogeneity rather than a direct estimate of stochasticity in observed biological time.

### 2.9 Identification of High-Entropy Individuals

To identify individuals exhibiting persistently high temporal instability, we defined a criterion based on the magnitude and temporal consistency of temporal entropy values. For each sample, the temporal entropy values were computed across all simulation steps, yielding a sequence *H*(*t*). Two summary statistics were derived for each sample: (i) the mean temporal entropy across all simulation steps and (ii) the proportion of steps exceeding a predefined threshold. Given a threshold *τ* = 0.6, the fraction of high-entropy steps was defined as

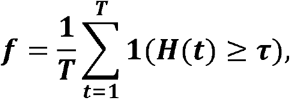

where *T* denotes the total number of simulation steps, and **1**(·) is the indicator function. Samples were classified as consistently high-entropy individuals if they satisfied both of the following criteria: (i) the mean temporal entropy across all steps was greater than or equal to *τ*, and (ii) the fraction of high-entropy steps was at least *f*_thr_, where *f*_thr_ = 0.8. This dual-criterion approach ensures that selected individuals exhibit elevated entropy on average and sustained temporal instability over the majority of the trajectory, thereby excluding individuals with only transient or sporadic fluctuations. The resulting subset represents a population with persistently unstable transcriptomic dynamics. The thresholds for and *τ* were *f* selected to balance sensitivity to instability and robustness against transient fluctuations.

### 2.10 Software and Implementation Details

All data preprocessing, statistical analyses, and machine learning procedures were performed using Python. Batch correction was conducted using the Scanpy Library (version 1.9.3), whereas classification models and cross-validation procedures were implemented with scikit-learn (version 1.3.0). Numerical integration of the inferred dynamic system and computation of temporal entropy were performed using custom Python scripts. PCA, smoothing spline fitting, and ridge regression were performed using standard scientific computing libraries, including NumPy (version 1.24.0) and SciPy (version 1.10.0). All analyses were conducted in a reproducible computational environment, and the parameter settings used in this study are available on request.

## 3 Results

### 3.1 Pairwise Classification Using Individual-Level Cross-Validation

First, we evaluated the feasibility of classifying psychiatric conditions based on blood transcriptomic profiles using pairwise classification across all diagnostic groups (CTRL, ADHD, ASD, and SCZ). The dataset comprised seven cohorts: two ADHD cohorts (44 and 32 samples), three ASD cohorts (104, 123, and 42 samples), and two SCZ cohorts (202 and 37 samples). Using individual-level stratified cross-validation, classification models were trained and evaluated for all six pairwise comparisons. The best-performing model for each pair, defined as the one with the highest AUROC among the three classifiers, is shown in Figure 1. Random forest models achieved high classification performance (AUROC > 0.8) for most pairs, except for CTRL vs. ASD. In contrast, linear models, including Elastic Net and linear SVM, exhibited performances near chance level for several pairs. These results suggest that blood transcriptome profiles contain sufficient information to discriminate between psychiatric conditions. However, the performance discrepancies across model types indicate that non-linear models may capture disease-related signals and cohort-specific or technical variations.

**Figure 1.**
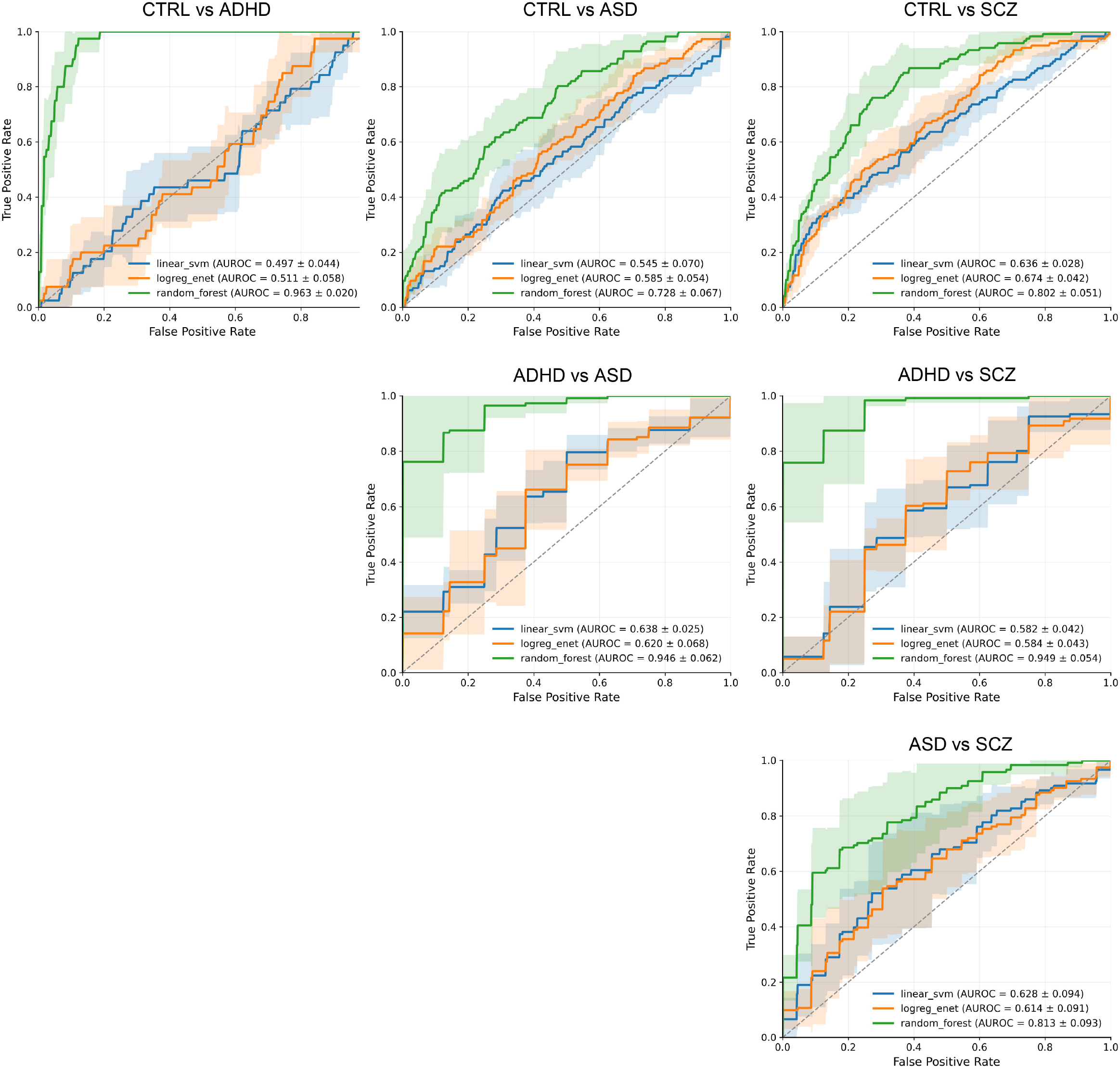
Classification performance of six diagnostic pairs based on individual-level cross-validation. Receiver operating characteristic (ROC) curves and area under the ROC (AUROC) values are shown for classification models trained on all six pairwise combinations of controls (CTRL), attention-deficit/hyperactivity disorder (ADHD), autism spectrum disorder (ASD), and schizophrenia (SCZ) using individual-level cross-validation. Green, orange, and blue lines represent random forest, Elastic Net–regularized logistic regression, and support vector machine (SVM) models, respectively. Shaded bands indicate the standard deviation of ROC curves across folds, calculated from fold-specific ROC curves.

### 3.2 Generalization Performance Using Study-Holdout Validation

To assess whether the observed classification performance reflected true disease-related signals or cohort-specific artifacts, we evaluated model performance using study-holdout cross-validation, in which the entire cohorts were excluded from training and used as independent test sets. Under this stringent evaluation framework, classification performance decreased substantially across all pairwise comparisons (Figure 2). For most pairs, AUROC values ranged from 0.5 to 0.7, and several comparisons, including CTRL vs. ASD and CTRL vs. ADHD, approached chance-level performance. Even for comparisons that retained modest discriminative abilities, such as CTRL vs. SCZ, ADHD vs. ASD, and ADHD vs. SCZ, AUROC values remained at approximately 0.6–0.7. This reduction in performance suggests that much of the predictive signal captured under individual-level cross-validation was driven by study-specific differences, such as variations in experimental platforms and cohort characteristics rather than robust disease-related biology. However, the residual performance above chance level in certain comparisons indicates that disease-related information is still partially encoded in blood transcriptomic profiles. Overall, these findings highlight the importance of evaluating generalizability across independent cohorts and suggest that conventional diagnostic labels may not fully align with the underlying biological structures captured by blood transcriptomic data.

**Figure 2.**
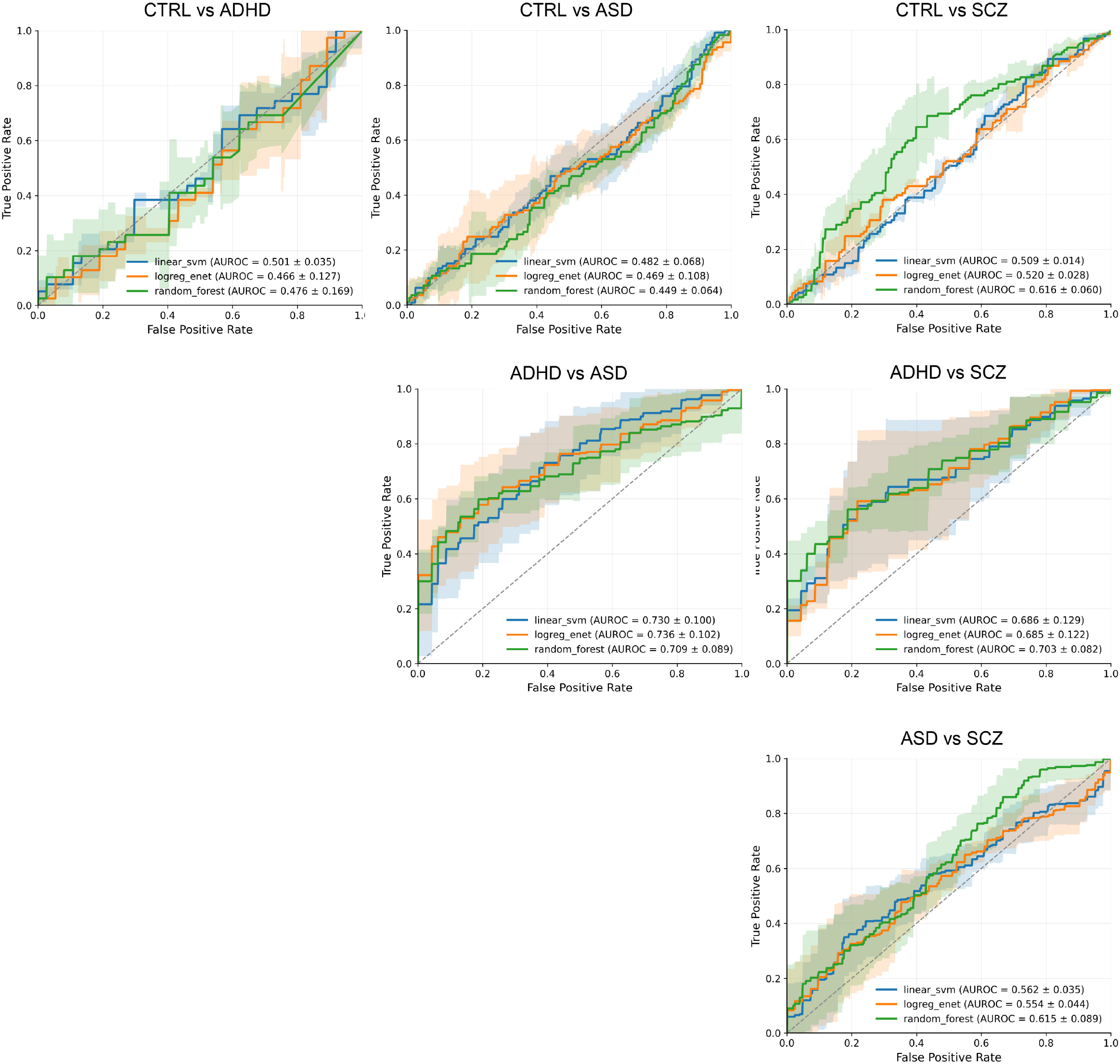
Classification performance of six diagnostic pairs based on study-holdout validation. ROC curves and AUROC values are shown for classification models trained on all six pairwise combinations of CTRL, ADHD, ASD, and SCZ using study-holdout validation. Green, orange, and blue lines represent random forest, Elastic Net–regularized logistic regression, and SVM models, respectively. Shaded bands indicate the standard deviation of ROC curves across folds, calculated from fold-specific ROC curves.

### 3.3 Attractor Analysis and Lack of Clear Diagnostic Clustering

To further investigate the relationship between transcriptomic structure and diagnostic labels, we performed attractor analysis based on the inferred dynamic system. We identified and visualized low-velocity regions corresponding to quasi-stable states and fixed points in principal component space (Figure 3). This analysis revealed fixed points and multiple low-velocity regions, indicating quasi-stable transcriptomic states. However, these regions did not correspond to distinct diagnostic categories. Instead, samples from various diagnostic groups (CTRL, ADHD, ASD, and SCZ) were broadly intermixed across the state space, including near the identified fixed points. This lack of clear clustering suggests that cross-sectional transcriptomic profiles do not segregate according to conventional diagnostic labels. These results highlight the limitations of interpreting cross-sectional data as static snapshots, which may obscure underlying dynamic transitions between states. Rather than forming discrete categories, the data appear to be organized along continuous and overlapping axes of variation. These findings support the view that psychiatric conditions are better understood as dynamically evolving systems in which observed transcriptomic differences reflect heterogeneous, time-varying biological processes not fully captured by current diagnostic classifications.

**Figure 3.**
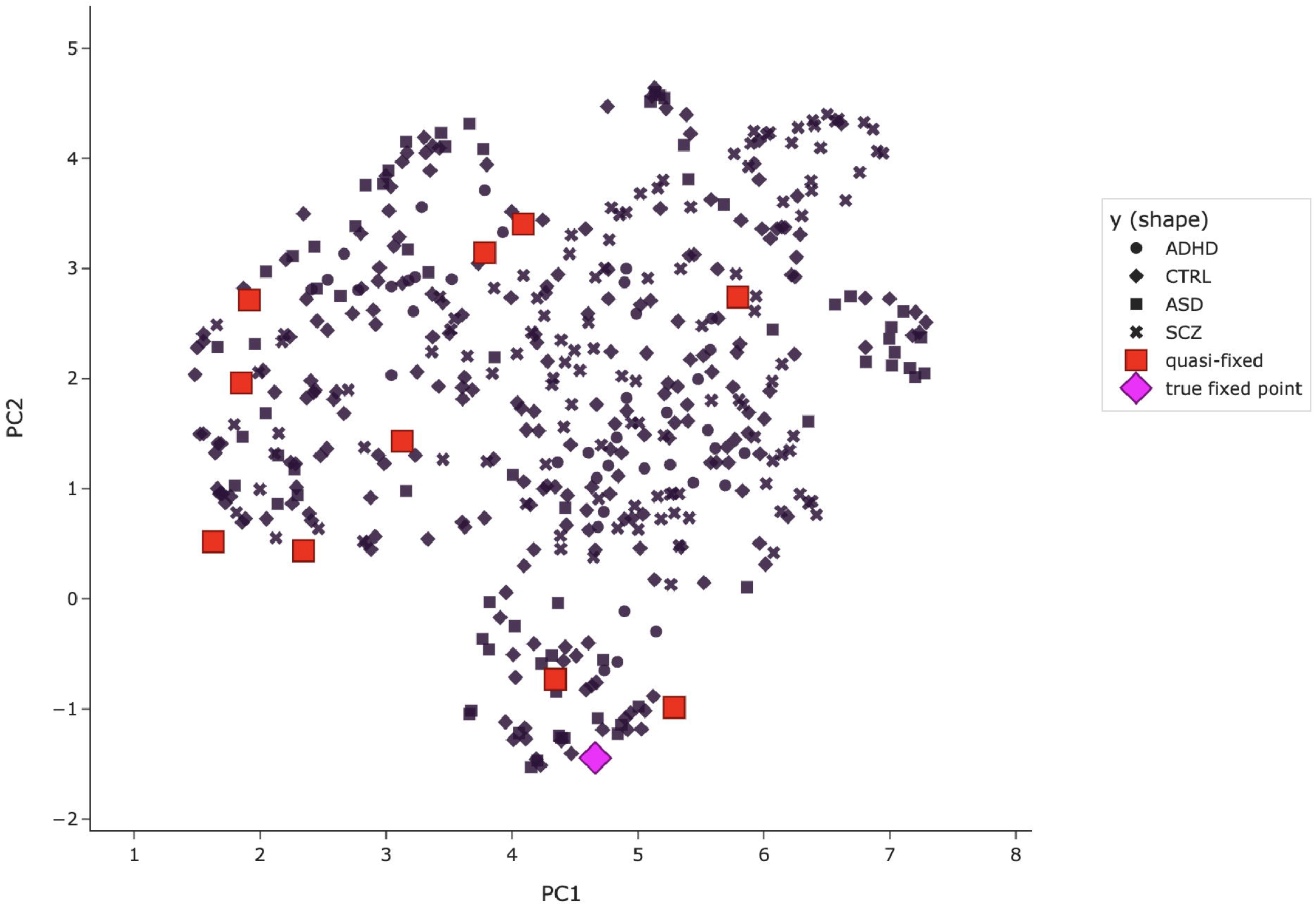
Attractor analysis based on transcriptomic profiles across all participants. All participants (CTRL, ADHD, ASD, and SCZ) are plotted in the principal component analysis (PCA) space. Marker shapes correspond to diagnostic groups as indicated in the legend. True and quasi-fixed points identified through attractor analysis are projected onto the PCA plane.

### 3.4 Temporal Entropy Reveals Dynamical Instability and Improves Classification

To capture the dynamic variability that is not apparent in static transcriptomic snapshots, we quantified temporal entropy for each individual based on inferred transcriptomic trajectories. The visualization of temporal entropy revealed that some individuals exhibited pronounced variability in state transitions over simulated time, indicating dynamically unstable transcriptomic behavior (Figure 4, Additional File 1). Such instability may reflect inconsistent or transitional biological states, lead to hypothesis that these individuals contribute to reduced classification performance. To test this, we excluded high-entropy individuals, as defined in the Methods section, and reevaluated classification performance using the study-holdout validation framework. Following the exclusion of high-entropy individuals, classification performance improved across most pairwise comparisons, with AUROC values exceeding 0.7 in all pairs, except for CTRL vs. ASD (Figure 5). In contrast, when classification models were trained and evaluated using only the excluded high-entropy individuals, performance consistently decreased compared to analyses of the remaining samples (Figure 6). These findings suggest that individuals characterized by high temporal entropy contribute disproportionately to reduced generalization performance. Furthermore, these results indicate that conventional diagnostic labels may include individuals with dynamically unstable states that are not captured by static classification models. By identifying and stratifying these individuals, temporal entropy enhances the robustness and generalizability of transcriptome-based psychiatric classification.

**Figure 4.**
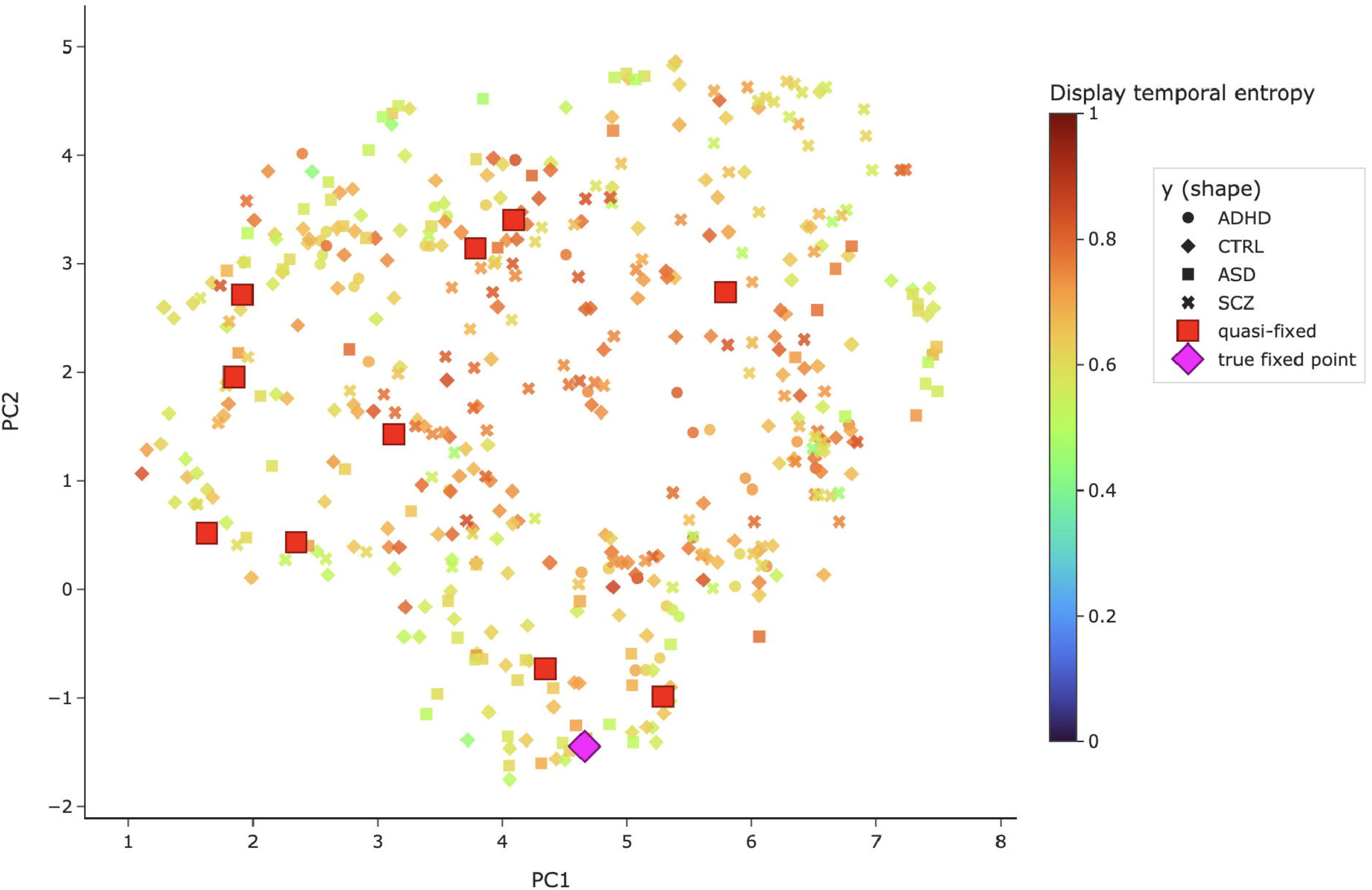
Visualization of temporal instability across participants based on temporal entropy. Snapshot at simulation step 100 of transcriptomic state evolution in PCA space, obtained by numerically integrating the inferred dynamic system *dX*/*dt* = *F*(*X*) derived from the attractor analysis (Figure 3). Marker shapes correspond to diagnostic groups (CTRL, ADHD, ASD, and SCZ), as indicated in the legend. Marker colors represent temporal entropy values for each participant. True and quasi-fixed points identified from the attractor analysis are projected onto the PCA plane.

**Figure 5:**
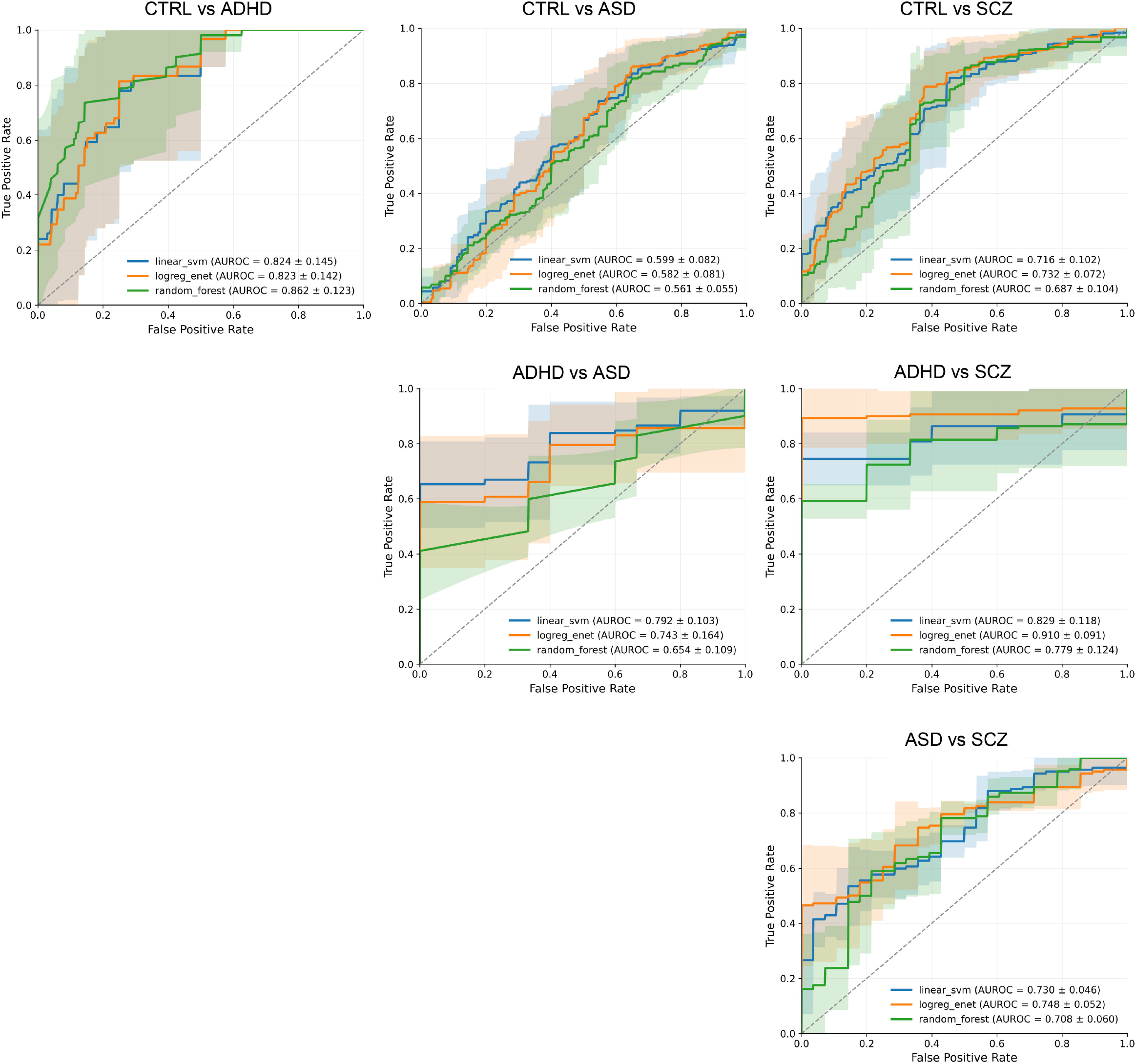
Classification performance after exclusion of high-entropy individuals. ROC curves and AUROC values are shown for classification models trained on all six pairwise combinations of CTRL, ADHD, ASD, and SCZ after excluding individuals with consistently high temporal entropy. Models were evaluated using study-holdout validation. Green, orange, and blue lines represent random forest, Elastic Net–regularized logistic regression, and support vector machine (SVM) models, respectively. Shaded bands indicate the standard deviation of ROC curves across validation splits, calculated from split-specific ROC curves.

**Figure 6:**
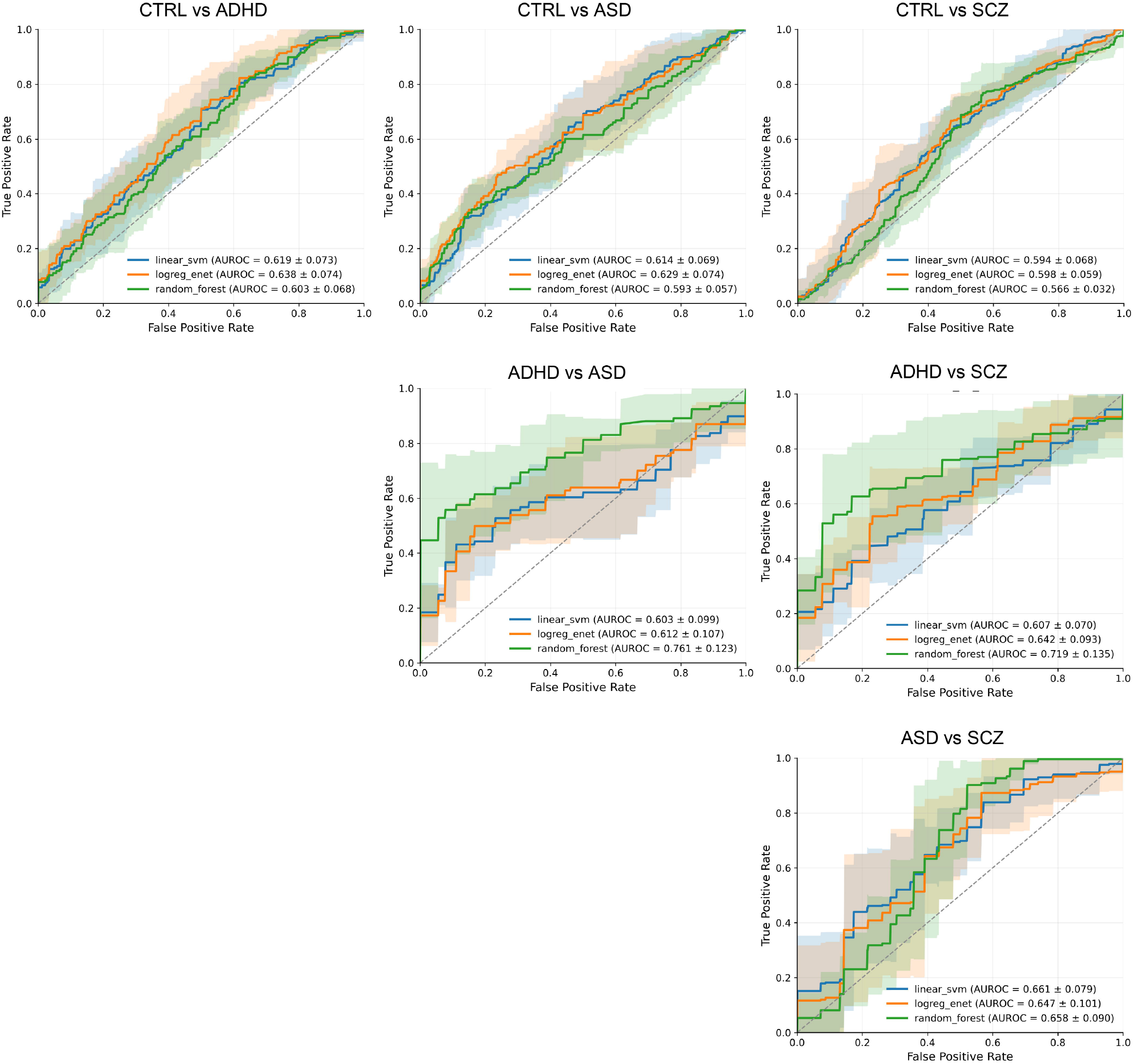
Classification performance using only high-entropy individuals. ROC curves and AUROC values are shown for classification models trained and evaluated on subsets consisting exclusively of individuals with consistently high temporal entropy. Models were evaluated using study-holdout validation. Green, orange, and blue lines represent random forest, Elastic Net–regularized logistic regression, and support vector machine (SVM) models, respectively. Shaded bands indicate the standard deviation of ROC curves across validation splits, calculated from split-specific ROC curves.

## 4 Discussion

This study investigated the feasibility and generalizability of psychiatric classification based on blood transcriptomic data from multiple independent cohorts. Three main findings were observed. First, high classification performance was achieved under individual-level cross-validation; however, this performance decreased significantly during study-level holdout validation, indicating limited cross-cohort generalizability. Second, attractor analysis revealed that transcriptomic states did not form distinct clusters corresponding to conventional diagnostic categories; instead, they exhibited overlapping and continuous structures in the state space. Third, stratification based on temporal entropy identified a subset of individuals with dynamically unstable transcriptomic trajectories, and excluding these individuals improved classification performance across most diagnostic pairs. Together, these findings suggest that variability in transcriptomic dynamics critically affects the inconsistency of classification performance and emphasizes the importance of incorporating dynamic features in developing biologically informed models of psychiatric disorders.

The first major finding of this study is the marked discrepancy between high classification performance in individual-level cross-validation and substantially reduced performance in study-level holdout validation. This result aligns with established concerns regarding batch effects and study-specific confounders in high-throughput transcriptomic data (9). Differences in experimental platforms, sample processing, and population characteristics can introduce systematic variations unrelated to disease status, which machine learning models may inadvertently exploit. When these confounding factors correlate with diagnostic labels, predictive models may learn study-specific patterns rather than biologically meaningful disease signals, leading to an overestimation of classification performance (10). The observation that non-linear models, particularly random forests, achieve higher performance than linear models under individual-level cross-validation further supports this interpretation, as these models capture complex, potentially non-biological patterns in the data. These findings do not imply that blood transcriptomic data lack disease-related information; rather, they indicate that a substantial portion of the apparent predictive signal reflects cohort-specific biases, underscoring the necessity for rigorous validation frameworks. In this context, study-level holdout validation provides a more realistic estimate of model performance in real-world scenarios in which models are applied to previously unseen populations.

The second major finding of this study was that attractor analysis did not reveal distinct clusters corresponding to conventional diagnostic categories but instead identified overlapping, continuous transcriptomic structures. This observation suggests that molecular variation in peripheral blood is not organized into discrete disease-specific states but reflects multiple underlying biological axes. These axes include immune activation, stress responses, developmental stages, medication effects, and cellular composition, which together shape the transcriptomic landscape. This interpretation aligns with previous studies, indicating that peripheral blood gene expression only partially reflects brain-specific pathology and is strongly influenced by systemic and cellular factors. Variations in immune cell composition have been shown to account for a significant portion of transcriptomic differences in psychiatric cohorts (18), and similar findings have been reported in ASD, where observed gene expression signatures are partly driven by differences in immune-related cell populations. From this perspective, the absence of clear clustering does not indicate a failure of transcriptomic approaches but reflects a mismatch between the multidimensional biological structure of the data and the categorical nature of current diagnostic systems. Additionally, the use of cross-sectional data limits the ability to resolve such structures because it captures only a static snapshot of inherently dynamic processes. Thus, the observed overlap across diagnostic groups may arise from biological heterogeneity and unobserved temporal transitions between states.

The third major finding of this study was that temporal entropy-based stratification improved the classification performance by identifying individuals with dynamically unstable transcriptomic trajectories. Excluding high-entropy individuals led to a consistent increase in classification performance across most diagnostic pairs, suggesting that these individuals disproportionately contributed to reduced generalizability. This finding underscores the importance of considering static molecular states and stability of the underlying biological dynamics. In contrast to conventional classification approaches that assume all samples can be assigned to a fixed category, the present findings support a two-step framework: (i) first assessing the stability or reliability of a sample and (ii) subsequently performing classification on a subset of individuals with more stable biological profiles. This approach aligns with emerging perspectives in clinical prediction modeling, emphasizing the need to account for uncertainty and define conditions under which predictions may be unreliable. Temporal entropy does not directly measure diagnostic correctness but captures a form of dynamic instability that may reflect biological heterogeneity, transitional states, or sensitivity to external perturbations. Clinically, individuals with high temporal entropy may require further evaluation or longitudinal monitoring rather than immediate categorical classification. Thus, temporal entropy may serve as a complementary biomarker, enhancing the robustness and interpretability of transcriptome-based psychiatric models by quantifying uncertainty in underlying biological states.

The concept of temporal entropy can be interpreted within the broader theoretical framework of biological robustness and resilience. In complex biological systems, robustness refers to the ability to maintain functional stability amid perturbations, whereas resilience describes the capacity to recover from disturbances and return to a stable state. These properties have been extensively studied in systems biology and ecological dynamics, where the loss of stability is often associated with increased variability and critical transitions between states (14,19). In mental health, similar concepts suggest that psychiatric symptoms may emerge when underlying regulatory systems lose stability and approach their tipping points (16). From this perspective, temporal entropy serves as a quantitative proxy for reduced robustness and resilience in transcriptomic dynamics. High temporal entropy indicates increased variability and incoherence in state transitions, suggesting that the system operates in a less stable regime and is more susceptible to perturbation. This interpretation extends beyond pathological states and applies to CTRL. Differences in temporal entropy may reflect variability in physiological stability, adaptability, or stress responsiveness, even among individuals without a clinical diagnosis. This broader applicability highlights the potential of temporal entropy as a general biomarker of system-level stability rather than a disease-specific indicator. In this framework, mental health can be viewed as the absence of disease and the presence of stable and resilient biological dynamics. Consequently, interventions such as pharmacological treatments, physical activity, and dietary modifications may promote stable system dynamics. Future research should investigate whether temporal entropy can be modulated through such interventions and whether reductions in entropy correlate with improved clinical or functional outcomes.

This study has several limitations. First, the inferred transcriptomic dynamics were based on pseudotime derived from cross-sectional data and did not represent true temporal processes. Although this approach enables approximation of latent trajectories, the estimated velocity reflects variations along the dominant axis of gene expression rather than actual time-dependent changes. This limitation relates to approaches such as pseudo-time analysis and RNA velocity in single-cell studies; however, their application to bulk transcriptomic data may be more susceptible to confounding factors, including differences in cell type composition and cohort-specific variation. Second, peripheral blood transcriptomic profiles are strongly influenced by cellular composition, environmental exposure, medication status, and other state-dependent factors. Although batch correction was applied to mitigate study-level differences, residual confounding may still affect inferred dynamics and temporal entropy estimates. Future studies incorporating cell-type deconvolution methods (18) or directly measuring cell composition data may clarify the biological interpretation of the observed patterns. Third, this study was based on publicly available datasets and represented a retrospective secondary analysis. Therefore, the proposed framework was not validated in independent prospective cohorts. The interpretation of high temporal entropy as a marker of instability or uncertainty remains indirect and cannot be assumed to correspond to diagnostic inaccuracy or clinical misclassification. Future research should address these limitations by (i) validating the proposed framework using independent datasets, (ii) incorporating longitudinal sampling to directly assess temporal stability, (iii) controlling for cell type composition and other confounding variables, and (iv) evaluating the clinical utility of temporal entropy in real-world decision-making contexts, including calibration and decision curve analysis.

## 5 Conclusions

This study demonstrated that blood transcriptomic data contain biologically relevant information for psychiatric classification. However, conventional evaluation approaches may overestimate performance because of cohort-specific effects. By adopting a dynamic systems perspective, we showed that transcriptomic states are better represented as continuous and overlapping structures than as discrete diagnostic categories. We introduced temporal entropy as a quantitative measure of dynamic instability, enabling the identification of individuals with less stable transcriptomic profiles that are less amenable to reliable classification. Stratification based on this metric improves cross-cohort generalizability and provides a framework for incorporating uncertainties into molecular prediction models. These findings suggest that integrating dynamic features and system-level stability into analytical frameworks may enhance the robustness and interpretability of psychiatric biomarkers. Future studies are warranted to validate this approach in longitudinal and clinically well-characterized cohorts and explore its potential utility in guiding personalized interventions.

## 6 List of Abbreviations

(ADHD): attention-deficit/hyperactivity disorder
(ASD): autism spectrum disorder
(SCZ): schizophrenia
(CTRL): healthy controls
(GEO): Gene Expression Omnibus
(SVM): support vector machine
(ROC): receiver operating characteristic
(OOF): out-of-fold
(TPR): true positive rate
(FPR): false positive rate
(AUROC): area under the ROC curve
(PC): principal component
(PCA): principal component analysis

## 7 Ethics approval and consent to participate

Not applicable.

## 8 Consent for publication

Not applicable.

## 9 Availability of data and materials

Not applicable.

## 10 Competing interests

The authors declare the following financial interests and personal relationships that may be considered potential competing interests.

TS is an employee of Rhelix Inc. RN is the founder and chief executive officer of this company.

## 11 Funding

The authors did not receive any funding for this study.

## 12 Authors’ contributions

Tatsuma Shoji (TS) performed the data analysis, generated all figures and tables, and drafted the manuscript. Ryo Nakaki (RN) led the project’s overall direction and management.

## 13 Acknowledgements

We thank Editage (www.editage.com) for their assistance with English language editing. The contributions of these individuals were indispensable for the successful execution of this study.

